# Long term maintenance of heterologous symbiont association in *Acropora palmata* on natural reefs

**DOI:** 10.1101/2022.08.13.503876

**Authors:** Holland Elder, Wyatt C. Million, Erich Bartels, Cory Krediet, Erinn Muller, Carly D. Kenkel

## Abstract

The sensitivity of reef-building coral to elevated temperature is a function of their symbiosis with dinoflagellate algae in the family Symbiodiniaceae. Changes in the composition of the endosymbiont community in response to thermal stress can increase coral thermal tolerance. Consequently, this mechanism is being investigated as a human-assisted intervention for rapid acclimation of coral in the face of climate change. Successful establishment of novel symbioses that increase coral thermal tolerance have been demonstrated in laboratory conditions; however, it is unclear how long these heterologous relationships persist in nature. Here, we show that *Acropora palmata* can form a novel symbiosis with *Durusdinium spp*. when reared in land-based aquaria. We tested the stability of this heterologous relationship by outplanting clonal replicates (ramets) of five *A. palmata* host genotypes to natural reefs in the lower Florida Keys. Amplicon sequencing analysis of ITS2-type profiles revealed that the majority of surviving ramets remained dominated by *Durusdinium spp*. two years after transplantation. However, 15% of ramets, including representatives of all genotypes, did exhibit some degree of symbiont shuffling or switching at six of eight sites, including complete takeover by site-specific strains of the native symbiont, *Symbiodinium fitti*. The predominant long-term stability of the novel symbiosis supports the potential effectiveness of symbiont modification as a management tool. However, the finding that 6–7 year-old coral can alter symbiont community composition in the absence of bleaching indicates that Symbiodiniaceae communities are indeed capable of greater flexibility under ambient conditions.

## Manuscript

Increasingly frequent and severe thermal stress events are the predominant threat to reef ecosystems in the Anthropocene. Thermal stress perturbs the photosynthetic mechanism and can result in loss or expulsion of the dinoflagellate algae in the process known as bleaching ^1^. While persistent bleaching can result in coral death due to the loss of autotrophic carbon contributed by the symbionts ^2^, bleaching can also precipitate an acclimatory change in the composition of the endosymbiont community that can subsequently increase coral thermal tolerance ^3,4^. For coral hosting multiple symbiont types, changes in their relative abundance, or ‘shuffling’, can facilitate survival and recovery ^3^. The ability for adult coral to acquire completely new symbiont types following bleaching was also recently confirmed ^5^. Although this capacity for flexibility is not universal ^6^, and is influenced by environmental cues and physiological trade-offs ^7^, symbiont shuffling and/or switching has been proposed as a key target for human-assisted evolution ^8^. Successful establishment of novel symbioses that increase coral bleaching tolerance have been demonstrated in laboratory conditions ^9^, however it is unclear how long these heterologous relationships persist, especially in natural reef environments. In a key test of the ultimate utility of this approach, we investigate the long-term stability of a heterologous coral symbiosis in the context of reef restoration.

*Acropora palmata* were once one of the dominant habitat builders of shallow Caribbean reefs ^10^. Unprecedented demographic declines since the mid-1970’s have resulted in extensive propagation efforts in both land and ocean-based nursery systems to both maintain existing genetic diversity and produce biomass for outplanting efforts aimed at restoring ecosystem structure and function ^11^. Naturally occurring populations of *A. palmata* are highly specific in their symbiosis with *Symbiodinium fitti*, formerly ITS2 type A3, ^12,13^. Single nucleotide polymorphism (SNP) genotyping of *ex situ* nursery-reared lines of *A. palmata*, however, revealed an almost exclusive association with *Durusdinium spp*. (Fig S1A, Supplementary Methods). This provided a unique opportunity to test the maintenance of this heterologous symbiotic relationship after outplanting back to the reef environment.

In April 2018, we transplanted three clonal replicates (ramets) of five exclusively *ex situ* nursery-reared *A. palmata* genotypes to nine reefs in the lower Florida Keys (n=27 ramets per genet) to evaluate the long-term stability of the *Durusdinium* association on natural reefs using periodic phenotypic monitoring and ITS2 amplicon sequencing for symbiont typing using the SymPortal pipeline ^14^, (Supplemental methods, Figure 1). A random sub-sampling of 10-12 ramets per genet confirmed that no *Symbiodinium fitti* were detectable prior to outplanting (T0, Supplemental methods, Figure 2A). Four genets were exclusively dominated by a *Durusdinium* ITS2 type, whereas genet 13-XK showed a consistent co-infection with a *Cladocopium* ITS2 type composed of three co-occurring C1 amplicon sequence variants (Figure 2A). A PCoA using Bray-Curtis dissimilarity of T0 samples confirmed a significant effect of genotype (p < 0.001) with 13-XK driving divergence (Figure 2B).

**Figure 1.**
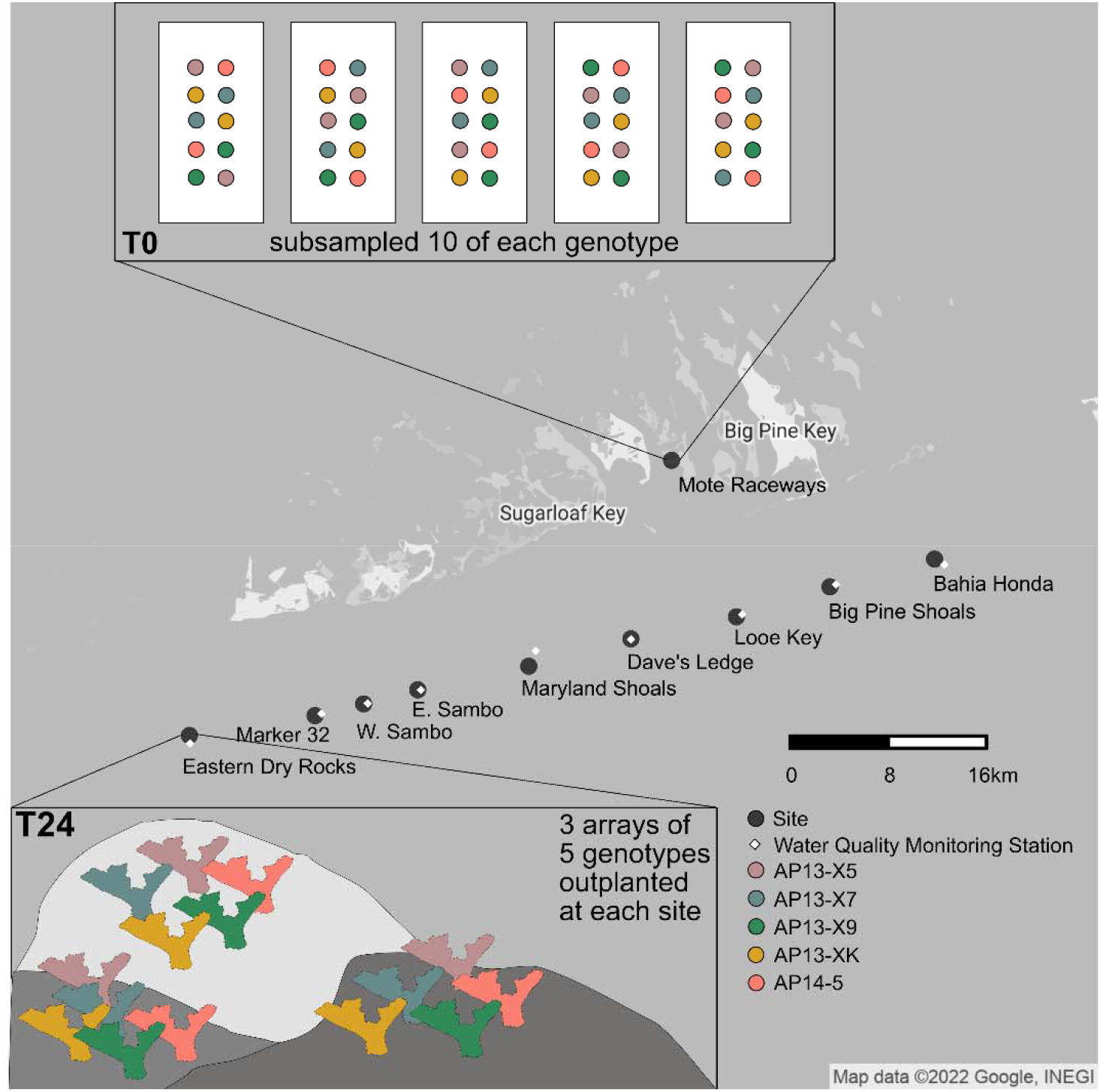
Experimental setup showing genotypes subsampled at time point 0 (T0), the locations of outplanting sites, the locations of water quality monitoring stations, and the genotype array strategy at each site. All surviving corals were sampled at time point 24 (T24).

**Figure 2.**
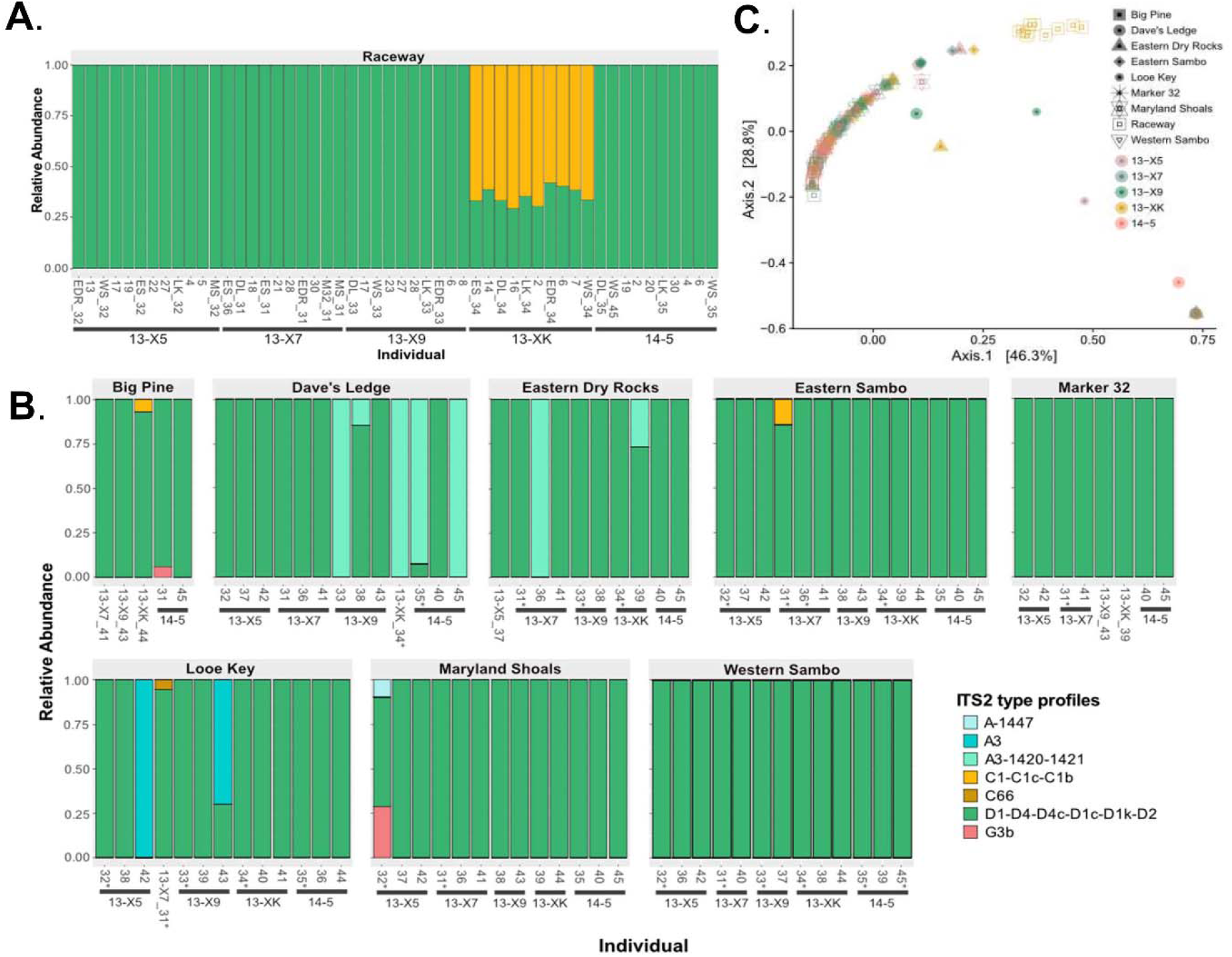
Relative abundance and PCoA plots showing that symbiont community did not significantly change after two years of outplanting. (**A.**) The relative abundance plot shows the ITS2 symbiont profiles of each individual coral grouped by genotype that was subsampled at T0. Each bar represents an individual and each is labeled with either their ex situ nursery frag label or their outplant site ID. Color represents symbiont type. (**B.**) The relative abundance plot shows the ITS2 symbiont profiles of each individual coral grouped by genotype that was sampled at T24. Each bar represents an individual. They are grouped by site, then genotype, and then frag id. Asterisks (*) identify individuals that were also sampled at T0. Color represents symbiont type. (**C.**) A PcoA plot that shows that symbiont profiles did not significantly change after two years of being outplanted. Color represents coral genotype and shape represents site. The ex situ nursery and T0 is included as a site.

No bleaching was observed during quarterly site visits, but site-specific mortality was evident, with 100% ramet mortality at Bahia Honda following 18 months (Supplemental Methods, Table S2). *Durusdinium* remained the dominant ITS2 profile after two years of outplanting (T24, Fig. 2B, C). Surviving co-infected 13-XK ramets also shuffled to *Durusdinium* dominance across sites (Fig. 2C) and a PERMANOVA confirmed genotype was no longer a significant factor (p = 0.764). Uptake of novel symbionts was detected, however, with site specific patterns (PERMANOVA, p < 0.031, Fig. 2C). Acquisition of the homologous symbiont, *Symbiodinium fitti* (A3) was observed at four sites, with ITS2 type profiles indicating potential sub-species or strain-level differences between Dave’s Ledge, Eastern Dry Rocks, Looe Key, and Maryland Shoals (Fig. 2C). Two ramets of 13-X7 acquired *Cladocopium* types, including one apparently novel strain (C66) although their relative abundance remained low (Figure 2C). A *Gerakladium* (G) type profile was also detected in ramets of two different genets at two different sites, also at low relative abundance (Fig. 2C).

The general long-term stability of the heterologous symbiosis between *A. palmata* and *Durusdinium spp*. on natural reefs supports the potential effectiveness of symbiont modification as a management intervention. Infection of coral larvae from mass spawning events with thermally tolerant symbionts and outplanting them to degraded reefs could also be a restoration tool ^15^. However, the occasional acquisition of homologous, site-specific strains of *Symbiodinium fitti* indicates that the temporal dynamics of symbiosis establishment in coral may be more complicated than previously thought. While corals can take up symbionts in their larval phase, the current consensus is that the majority of symbionts are taken up after settlement. New recruits and juvenile corals can form a variety of novel symbiont associations ^16^, but this diversity is generally pruned to one numerically dominant taxon per genus over the first few years of life ^17^. Subsequent changes to the dominant taxon occur in response to bleaching-level stress ^5,18,19^.

The *A. palmata* lines used in this study were produced using individual larvae from batch crosses in 2013/14. Although physically small, at the time of outplanting, experimental ramets were 4-5 years old, which under normal field conditions is the age of sexual maturity for this species ^20^. The finding that adult corals can switch their symbionts *in situ* in the absence of bleaching suggests that acquisition and winnowing may be size-dependent rather than age-dependent process. It is also possible that Symbiodiniaceae communities are indeed capable of greater flexibility under ambient conditions ^21^. However, the stability of the heterologous symbiosis in most individuals suggests that infection with a novel symbiont type as a recruit and maintenance till maturity could produce a stable symbiosis.

## Supporting information

Supplemental Methods and Results

## Acknowledgements

We thank the staff of Mote Marine Labs for their assistance in producing and maintaining the coral genotypes used in this study. We also thank all graduate students of the Cnidarian Evolutionary Ecology Lab and Mote Marine Labs staff who participated in monitoring of these corals during the outplant period.

## Code and Data Availability

Sequence Access: NCBI, Accession: PRJNA868513 https://github.com/hollandelder/OutplantedA.palmITS2

## Authors’ contributions

C.D.K., E.M., C.K, W.M., and H.E. conceived the experiments and analysis. W.M., E.B., C.K., and C.D.K. conducted experiments. E.M. and E.B. provided advice and logistical support through the experiment. H.E. conducted genetic and statistical analysis and wrote the manuscript. C.D.K. and E.M. provided advice and support for analysis. C.D.K., E.M. and C.K. helped write the manuscript.

