## Supplemental Methods and Results for "Long term maintenance of heterologous symbiont association in *Acropora palmata* on natural reefs"

**Supplemental Text**

**Supplemental Methods**

*Experimental Setup and Samples Gathered*

Individual larvae representing five unique genotypes of *Acropora palmata* (Figure 1) from multi-parent batch crosses produced in 2013 (denoted with prefix ‘13’) and 2014 (denoted with prefix ‘14’) were settled on aragonite plugs and grown out in Mote Marine Lab’s International Center for Coral Reef Research and Restoration (IC2R3), *ex situ* raceways supplied with flow through seawater. Temperatures ranged from 24-25̊C in the winter to 26-28̊C in the summer. Also, pH ranged from 7.7-7.9. Salinity ranged from 37-39 ppt. Ramets were regularly rotated every month to minimize tank effects. Over time, once sufficient growth was achieved, genotypes were microfragmented ^1^ to produce clonal replicates (ramets). In April of 2018, 27 replicate ramets of each genotype (N=135 total) were identified for outplanting and small tissue samples were taken before outplanting from ten random ramets of each genotype except for 13-X5 of which 12 ramets were sampled to determine symbiont community composition of each genotype prior to outplanting (T0). Samples were flash frozen in liquid nitrogen and stored at -80 ̊ C. Ramets were then outplanted in triplicate arrays at nine field sites in the lower Florida Keys (Supplemental Table 1, Supplemental Figure 1). These sites were chosen because they are active restoration sites where the corals would be in proximity to other corals in symbiosis with *Symbiodinium fitti* ^2,3^, which gave them the opportunity to take up their wild type symbiont. Of those individuals that were subsampled at T0, 5-6 of them were placed at each site, so that they could be resampled at time point 24 months (T24). Coral outplants were monitored every three months to assess performance. No bleaching was observed throughout the course of the experiment ^4^, but some differential mortality was evident (Supplemental Table 2). Following 24 months, tissue samples were taken from each surviving ramet (T24, N= 92 total ramets), flash frozen within 5 minutes of collection and stored at -80 ̊ C.

*Amplicon Library Preparation*

Tissue was removed from the skeleton and homogenized in Aurum Total RNA lysis buffer using the OMNI International bead rupture elite with sterile glass beads. DNA was extracted from all samples using the BioRad Aurum Total RNA Mini Kit. The SYM_VAR_5.8S2 (forward: 5’-GTGACCTATGAACTCAGGAGTCGAATTGCAGAACTCCGTGAACC-3’) and SYM_VAR_REV (reverse: 5’-CTGAGACTTGCACATCGCAGCCGGGTTCWCTTGTYTGACTTCATGC-3’) primers were used to amplify the ITS2 locus following the protocol described in ^5^. PCR reactions contained: 0.5µl of equimolar (10mM) dNTPs, 5 µl 5X Q5 PCR buffer (New England Bio Labs), 0.25 µl Q5 DNA polymerase (New England Bio Labs), 5 µl DNA (20ng/µl) template, 0.25 µl of each primer (10 µM), 0.25 µl BSA (20 mg/µl) (New England Bio Labs), and 13.5 µl Milli-Q ultrapure water per reaction. The amplification profile contained an initial denaturation step of 98°C for 30 seconds with the cycle starting at 98°C for 10 s, 56°C for 1 minute, and 72°C for 30 s each cycle with a final step 72°C for 5 min to finish. Samples amplified at 18 cycles of PCR. Three samples failed to amplify and were discarded from subsequent steps. Water was used as a negative control during PCR and no bands were observed. In a second four cycle PCR, Illumina primers with custom dual index barcodes were incorporated. Barcoding PCR reactions contained: 0.5µl of equimolar (10mM) dNTPs, 5 µl 5X Q5 PCR buffer (New England Bio Labs), 0.25 µl Q5 DNA polymerase (New England Bio Labs), 3 µl DNA template, 1 µl of each of two barcodes (10 µM), 0.25 µl BSA (20 mg/µl) (New England Bio Labs), and 14 µl Milli-Q ultrapure water per reaction. The amplification profile for barcoding contained an initial denaturation step of 98°C for 30 seconds with the cycle starting at 98°C for 10 s, 59°C for 1 minute, and 72°C for 30 s each cycle with a final step 72°C for 2 min to finish. Concentrations of each amplicon were determined using Picogreen (Thermo Fisher Scientific) and then pooled in equimolar amounts. The amplicon library was sequenced at University of Southern California Keck Molecular Genomics Core on the Illumina MiSeq V2 Nano sequencing platform (PE 250). Raw sequences from this study can be found at (BioProject: PRJNA868513).

*Analysis*

To identify ITS2 type profiles, sequences were analyzed using the Symportal desktop platform as in ^6^. Code can be accessed at (https://github.com/hollandelder/OutplantedA.palmITS2). SymPortal analysis determined ITS2 profiles from the presence and abundance ITS2 sequences or defining intragenomic variants (DIVS) across samples ^6^. SymPortal analysis calculated relative abundance ^6^. We did a principal coordinate analysis (PCoA) using Bray-Curtis dissimilarity on the ITS2 type profiles in the Phyloseq package ^7^. We used PERMANOVA tests (999 permutations) in the vegan software package ^8^ to evaluate sample dissimilarity between samples, between genotypes, between sites, and between time points.

**Supplemental Results**

After quality filtering, the mean number of reads per sample was 4137 reads retaining 95% of the original reads. The mean percentage of reads that mapped in the database was 90% with 3722+/-1074 ITS2 counts per sample.

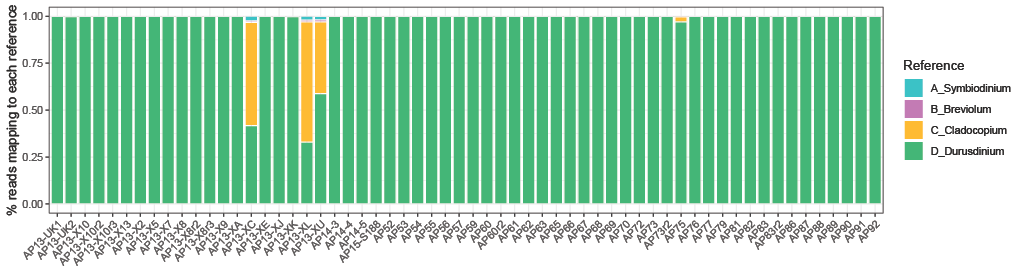

**Supplemental Figure 1**: Percent of reads that map to Symbiodiniaceae references for Mote Marine Lab *Acropora palmata* genotypes identified through 2bRAD SNP genotyping. Each bar represents an individual *A. palmata* genotype and color represents symbiont types.

**Supplemental Table 1**: The latitude and longitude of the

outplant sites and the water quality (WQ) monitoring stations

| **Outplant Site or WQ Station** | **Latitude** | **Longitude** |
| --- | --- | --- |
| Bahia Honda | 24.58908 | -81.242 |
| Big Pine Shoals | 24.56867 | -81.3266 |
| Looe Key | 24.54667 | -81.4021 |
| Dave's Ledge | 24.53051 | -81.4873 |
| Maryland Shoals | 24.51036 | -81.5699 |
| E. Sambo | 24.49305 | -81.6596 |
| W. Sambo | 24.48268 | -81.7033 |
| Marker 32 | 24.47408 | -81.7426 |
| Eastern Dry Rocks | 24.45948 | -81.8441 |
| Bahia Honda WQ | 24.585 | -81.2341 |
| Big Pine WQ | 24.5704 | -81.3217 |
| Looe Key WQ | 24.5484 | -81.3974 |
| Maryland Shoals WQ | 24.5216 | -81.5643 |
| E. Sambo WQ | 24.4929 | -81.6571 |
| Western Sambo WQ | 24.483 | -81.7 |
| Eastern Dry Rocks WQ | 24.4536 | -81.8437 |

**Supplementary Table 2**: Surviving coral numbers from for each genotype at each site

| **Site** | **Genotype** | **T0** | **T3** | **T6** | **T9** | **T12** | **T15** | **T18** | **T24** |
| --- | --- | --- | --- | --- | --- | --- | --- | --- | --- |
| Bahia Honda | AP13-X5 | 3 | 3 | 3 | 3 | 1 | 1 | 0 | 0 |
| Bahia Honda | AP13-X7 | 3 | 3 | 3 | 3 | 0 | 0 | 0 | 0 |
| Bahia Honda | AP13-X9 | 3 | 3 | 3 | 3 | 3 | 3 | 0 | 0 |
| Bahia Honda | AP13-XK | 3 | 3 | 3 | 2 | 2 | 1 | 0 | 0 |
| Bahia Honda | AP14-5 | 3 | 2 | 1 | 1 | 0 | 0 | 0 | 0 |
| Big Pine | AP13-X5 | 3 | 3 | 3 | 3 | 3 | 3 | 3 | 1 |
| Big Pine | AP13-X7 | 3 | 3 | 3 | 3 | 3 | 3 | 3 | 1 |
| Big Pine | AP13-X9 | 3 | 3 | 3 | 3 | 3 | 3 | 3 | 1 |
| Big Pine | AP13-XK | 3 | 3 | 3 | 3 | 3 | 3 | 3 | 1 |
| Big Pine | AP14-5 | 3 | 3 | 3 | 3 | 3 | 3 | 3 | 1 |
| Dave's Ledge | AP13-X5 | 3 | 3 | 3 | 3 | 3 | 3 | 3 | 3 |
| Dave's Ledge | AP13-X7 | 3 | 3 | 3 | 3 | 3 | 3 | 3 | 3 |
| Dave's Ledge | AP13-X9 | 3 | 3 | 3 | 3 | 3 | 3 | 3 | 3 |
| Dave's Ledge | AP13-XK | 3 | 3 | 2 | 2 | 2 | 2 | 2 | 2 |
| Dave's Ledge | AP14-5 | 3 | 3 | 3 | 3 | 3 | 3 | 3 | 3 |
| Eastern Sambo | AP13-X5 | 3 | 3 | 3 | 3 | 3 | 3 | 3 | 3 |
| Eastern Sambo | AP13-X7 | 3 | 3 | 3 | 3 | 3 | 3 | 3 | 3 |
| Eastern Sambo | AP13-X9 | 3 | 3 | 3 | 3 | 3 | 3 | 3 | 3 |
| Eastern Sambo | AP13-XK | 3 | 3 | 3 | 3 | 3 | 3 | 3 | 3 |
| Eastern Sambo | AP14-5 | 3 | 3 | 3 | 3 | 3 | 3 | 3 | 3 |
| Eastern Dry Rocks | AP13-X5 | 3 | 2 | 1 | 1 | 1 | 1 | 1 | 1 |
| Eastern Dry Rocks | AP13-X7 | 3 | 3 | 3 | 3 | 3 | 3 | 3 | 3 |
| Eastern Dry Rocks | AP13-X9 | 3 | 2 | 2 | 2 | 2 | 2 | 2 | 2 |
| Eastern Dry Rocks | AP13-XK | 3 | 2 | 2 | 2 | 2 | 2 | 2 | 2 |
| Eastern Dry Rocks | AP14-5 | 3 | 2 | 2 | 2 | 2 | 2 | 2 | 2 |
| Looe Key | AP13-X5 | 3 | 3 | 3 | 3 | 3 | 3 | 3 | 3 |
| Looe Key | AP13-X7 | 3 | 3 | 2 | 2 | 2 | 2 | 2 | 2 |
| Looe Key | AP13-X9 | 3 | 3 | 3 | 3 | 3 | 3 | 3 | 3 |
| Looe Key | AP13-XK | 3 | 3 | 3 | 3 | 3 | 3 | 3 | 3 |
| Looe Key | AP14-5 | 3 | 2 | 2 | 2 | 2 | 2 | 2 | 2 |
| Marker 32 | AP13-X5 | 3 | 3 | 3 | 3 | 3 | 3 | 3 | 2 |
| Marker 32 | AP13-X7 | 3 | 2 | 2 | 2 | 2 | 2 | 2 | 2 |
| Marker 32 | AP13-X9 | 3 | 3 | 3 | 3 | 3 | 3 | 2 | 1 |
| Marker 32 | AP13-XK | 3 | 3 | 3 | 3 | 2 | 2 | 2 | 2 |
| Marker 32 | AP14-5 | 3 | 3 | 3 | 3 | 3 | 3 | 2 | 2 |
| Maryland Shoals | AP13-X5 | 3 | 3 | 3 | 3 | 3 | 3 | 3 | 3 |
| Maryland Shoals | AP13-X7 | 3 | 3 | 3 | 3 | 3 | 2 | 2 | 2 |
| Maryland Shoals | AP13-X9 | 3 | 3 | 3 | 3 | 3 | 3 | 3 | 2 |
| Maryland Shoals | AP13-XK | 3 | 3 | 3 | 3 | 2 | 2 | 2 | 2 |
| Maryland Shoals | AP14-5 | 3 | 3 | 3 | 3 | 3 | 3 | 3 | 3 |
| Western Sambo | AP13-X5 | 3 | 3 | 3 | 3 | 3 | 3 | 3 | 3 |
| Western Sambo | AP13-X7 | 3 | 3 | 3 | 3 | 3 | 3 | 3 | 3 |
| **Site** | **Genotype** | **T0** | **T3** | **T6** | **T9** | **T12** | **T15** | **T18** | **T24** |
| Western Sambo | AP13-X9 | 3 | 3 | 2 | 2 | 2 | 2 | 2 | 2 |
| Western Sambo | AP13-XK | 3 | 3 | 3 | 3 | 3 | 3 | 3 | 3 |
| Western Sambo | AP14-5 | 3 | 3 | 3 | 3 | 3 | 3 | 3 | 3 |
